# A Matched-filter Based Algorithm for Subcellular Classification of T-system in Cardiac Tissues

**DOI:** 10.1101/371328

**Authors:** Dylan F Colli, S Ryan Blood, Aparna C Sankarankutty, Frank B Sachse, Michael Frisk, William E Louch, Peter M Kekenes-Huskey

## Abstract

In mammalian ventricular cardiomyocytes, invaginations of the surface membrane form the transverse tubular system (T-system) which consists of transverse tubules (TTs) that align with sarcomeres and Z-lines as well as longitudinal tubules (LTs) that are present between Z-lines in some species. In many cardiac disease etiologies the T-system is perturbed, which is believed to promote spatially heterogeneous, dyssynchronous Ca^2+^ release and inefficient contraction. In general, T-system characterization approaches have been directed primarily at isolated cells and do not detect subcellular T-system heterogeneity. Here we present MatchedMyo, a matched-filter based algorithm for subcellular T-system characterization in isolated cardiomyocytes and millimeter-scale myocardial sections. The algorithm utilizes “filters” representative of TTs, LTs, and T-system absence. Application of the algorithm to cardiomyocytes isolated from rat disease models of myocardial infarction (MI), dilated cardiomyopathy induced via aortic banding (AB), and sham surgery confirmed and quantified heterogeneous T-system structure and remodeling. Cardiomyocytes from post-MI hearts exhibited increasing T-system disarray as proximity to the infarct increased. We found significant (p<0.05, Welch’s t-test) increases in LT density within cardiomyocytes proximal to the infarct (12±3%, data reported as mean ± SD, n=3) vs. sham (4±2%, n=5), but not distal to the infarct (7±1%, n=3). The algorithm also detected decreases in TTs within 5° of the myocyte minor axis for isolated AB (36±9%, n=3) and MI cardiomyocytes located intermediate (37±4%, n=3) and proximal (34±4%, n=3) to the infarct vs. sham (57±12%, n=5). Application of bootstrapping to rabbit MI tissue revealed distal sections comprised 18.9±1.0% TTs while proximal sections comprised 10.1±0.8% TTs (p<0.05), a 46.6% decrease. The matched filter approach therefore provides a robust and scalable technique for T-system characterization from isolated cells through millimeter-scale myocardial sections.

## INTRODUCTION

In ventricular cardiomyocytes, Ca^2+^-induced-Ca^2+^-release (CICR) begins with the propagation of an action potential along the cell membrane, eliciting sarcolemmal Ca^2+^ influx via L-type Ca^2+^ channels (LCCs). These channels are primarily localized within sarcolemma (SL) invaginations comprising the T-system[1]. Localized Ca^2+^ entry via LCCs triggers neighboring ryanodine receptors (RyRs) located in the sarcoplasmic reticulum (SR) membrane to open and release Ca^2+^ from SR stores [1–4]. The spatiotemporal synchrony and homogeneity of CICR is critical in maintaining efficacious contraction of the heart. In rabbit, canine, and human ventricular cardiomyocytes this transverse tubular system (T-system) is comprised predominantly of TTs coinciding with the Z-lines. In rodent ventricular cardiomyocytes, the T-system includes both TTs as well as significant longitudinal tubule (LT) and axial tubule components [5, 6]. In failing hearts, severe morphological changes in the T-system of ventricular cardiomyocytes promotes impairment of critical excitation-contraction coupling (ECC) Ca^2+^ handling mechanisms [1, 7] that drive Ca^2+^ dysregulation and cardiac dysfunction [8–10].

Advances in fluorescence-based microscopy offer outstanding potential to probe mechanisms governing T-system remodeling. However, the nature of T-system remodeling is diverse, with some pathologies almost strictly presenting tubule absence (TA), while others can exhibit tubules in non-transverse arrangements. Among the disease models that manifest evident changes in cardiomyocyte T-system, distinctive patterns have emerged from reduced TT density in type-II diabetic mice [11] to the coupled observation of TA and increased LT density with differing patterns for congenital heart failure [12], spontaneous hypertension/heart failure [13], and other disorders [14]. Substantial amounts of sub-micron resolution confocal microscopic images exist for several models of heart failure (concentric hypertrophy and dilated hypertrophy, for instance) in species including mice [9, 12, 15–17], rabbit [18–20], canine [21, 22], and human [23, 24]. However, quantitative assessments of subcellular T-system remodeling have been limited.

Characterization of images and correlating structural features with cardiac function [25] represent significant challenges that permit computational solutions. Several methods were introduced to automate the characterization of cardiomyocyte T-system. Among these approaches is the ‘TT index’ or ‘*TT_Power_*’, whereby the relative spacing and angle of TT arrangements are estimated by Fourier analysis. [26–28] *TT_Power_* correlates well with the severity of remodeling, but provides limited structural classification. More recent methods apply some level of preprocessing in order to essentially increase the amplitude of the signal arising from the TTs [26, 27]. The most common metric appears to be binary thresholding, which increases contrast by converting all pixels to binary values, depending on their value relative to a user-determined threshold. Others utilize morphological transformations which produce ‘skeletonized’ representations of the T-system which enhance the TT index. [26, 27, 29] Recent uses of techniques such as histogram equalization [30] have been tuned to improve existing analysis approaches [26, 27, 29]. The advantage of these approaches is that they assume little prior knowledge about the structure of the T-system, yet their applications have been limited to single cardiomyocyte preparations.

Here we present MatchedMyo, a matched-filter based image processing algorithm for the subcellular characterization of cardiomyocyte T-system. The method relies on images, or filters, representing features of interest, such as native TTs, LTs, and TA. In contrast to methods that binarize images to highlight features, this algorithm preserves a dynamic range of the original image. We outline the corresponding workflow in Fig. 1, which we use to characterize the total cellular content of intact TT structure, LTs, and TA in isolated sham, MI, and dilated cardiomyopathy via ascending aortic banding (AB) cells as well as MI tissue sections (cell and tissue preparation summarized in Sect. S.3.1).

**Figure 1:**
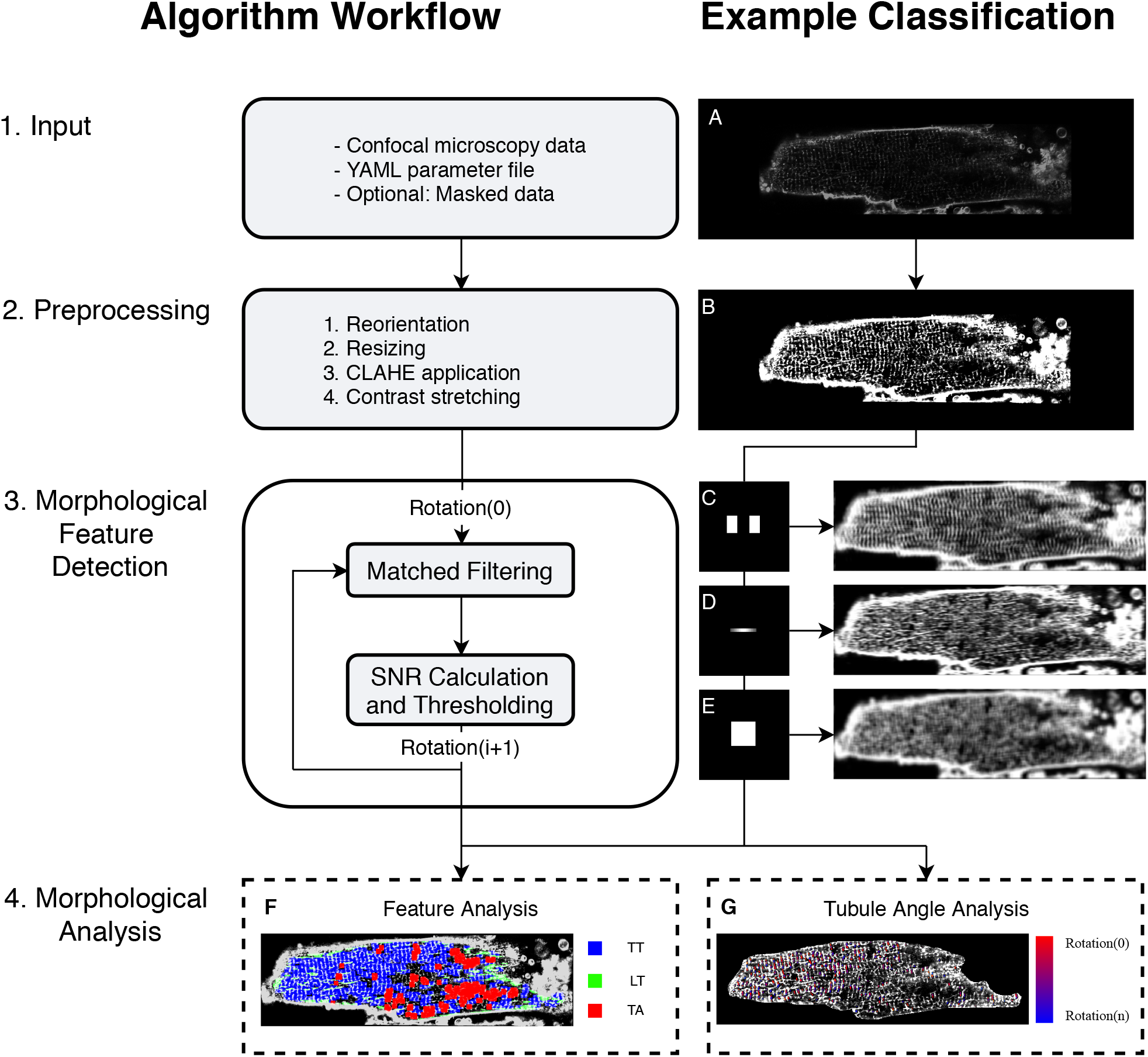
Unicellular matched filtering protocol. (1) Input of confocal microscopy data, YAML parameter file, and optional user generated mask for that data, where (A) shows sample input image.. YAML parameter file construction is explained in Sect. S.1.4. (2) Integrated preprocessing routines where (B) shows sample preprocessed image. (3) Morphological feature detection via matched filtering and signal-to-noise ratio calculation. (C-E) Matched-filters and sample signal-to-noise ratio calculation via convolution for TT, LT, and TA filters, respectively. (4) Morphological analysis of features, feature density, and feature rotation. (F) Sample morphological feature content analysis exhibiting high TT content relative to other morphological features. (G) Sample tubule striation angle analysis with high angle variance.

## MATERIALS AND METHODS

### Matched filter theory and application

We developed a computer program that analyzes pictures of cardiac cells and measures where structures involved in calcium handling are located. This is important because changes in cell calcium can contribute to disease. Our program was based on techniques from signal processing theory and used a technique called ‘matched filtering.’ In essence, our matched filtering approach scans cardiomyocytes for regions that resemble a representative image, or filter, of transverse and longitudinal tubule features. This scanning is accomplished by a mathematical operation, convolution, of the cardiomyocyte image with the TT filter. The result of this convolution is a new image, for which each pixel represents the probability of finding the filter at the corresponding location in the cardiomyocyte. We utilize a bank of rotated TT filters to tolerate moderate variations in TT orientation, followed by automated, post-processing techniques to classify detected features.

More formally, matched filtering determines whether a known signal, *s*, or in vector form, *ŝ*, is embedded in statistically uncorrelated noise, 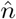, via the relationship

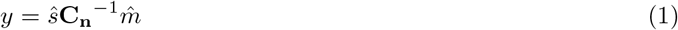

where *y* is the likelihood the known signal is embedded in the measured signal, 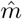 is the measured signal 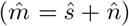, [31] and **C_n_** is the noise covariance 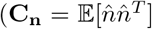, where 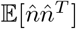 signifies the expectation of a random, spatially distributed noise vector, 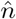). The overarching principle in matched filtering is to identify a matched filter, 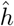, that maximizes the signal-to-noise ratio for a measurement, 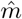, [31] via

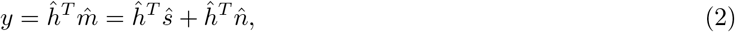

where 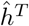 denotes the filter’s transpose. We utilize this approach in the first stage of our workflow in order to assess the likelihood that a filter representing a feature of interest is present in a given data set. Often the signal, or multiple instances thereof, is embedded within a larger data set 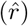, such as an image. In which case, determining the location of *ŝ* within 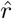 is commonly performed by convolving the filter 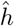 with the image, 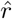, via

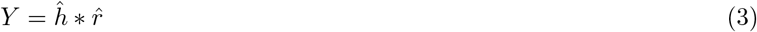

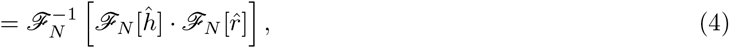

where *Y*(*x, y*) is the likelihood of finding the known signal, *ŝ* in the image, 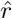 at (*x, y*), while 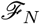 and 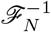 represent the discrete Fourier transform and inverse Fourier transform, respectively, to solve Eq. 3 via the convolution theorem. Probable detections of the filter 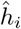 within *Y_i_* are then based on identifying positions at which the response is above a user-specified threshold criterion, *λ_i_*. To classify where cellular microstructure resembles known filters, we evaluate

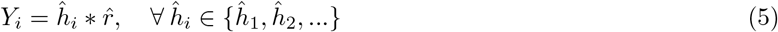

The optimal filter is chosen as max({*Y*_1_, *Y*_2_,…}), so long as there exists at least one *Y_i_* for which *Y_i_*(*x, y*) > *λ_i_*. Regions of the test confocal data that returned responses below the threshold parameters for all filters considered are designated as ‘uncharacterized.’ Matched filter data reported herein for TT, LT, and TA filters are reported as percent characterized cell area of total cell area for each filter.

#### Detection by convolution

Beyond classifying whether cellular microstructure resembles known filters, our workflow is tasked with designating regions of T-system absence as well as unclassified TT structure. For this reason, we present the following detection scheme. We assess the likelihood of subcellular content via

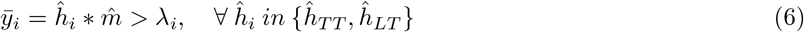

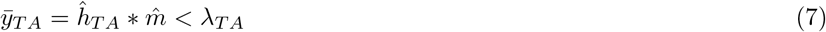

where * represents the convolution of the filter, 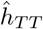, 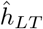, and 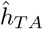, with the confocal microscopy image, 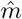, and *λ_i_* represents the user-determined threshold. A region is considered uncharacterized if no 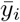 is indicated as a hit via this criterion.

#### Filter bank

In our approach, we additionally consider filter rotations to detect alternate tubule orientations through defining a bank of filter rotations for each matched filter spaced at 5 degree increments. The optimal rotation for a given filter is determined by evaluating each rotated filter with the data; the rotated filter that generates the highest above-threshold response is elected as the most likely orientation of a given feature, *j*:

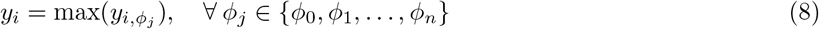

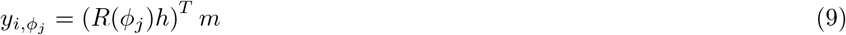

where *R*(*ϕ_j_*) represents the rotation of filter *h* by *ϕ_j_*.

#### Matched filter determination

Here we determine the matched filters based on manually identifying representative T-system structure in the confocal data. The three morphological characteristics that we identified were TT, LT, and TA regions. The filters are shown in Fig. 1. The TT filter consisted of two separated binary TT structural elements. The complement to this filter was utilized as a penalty filter as described in Sect. S.2.3. The LT filter consisted of a one sarcomere wide longitudinal structural element with varying intensity according to a normal distribution along the x axis. This allowed for greater specificity of LT versus TT detection by attenuating the spectral overlap of the filters. The TA filter was a sarcomere-length square smoothing filter. The presence of sub-threshold pixels in the smoothed image indicated the presence of sarcomere-sized regions of low signal density. Details for selecting optimal threshold parameters and corresponding receiver operator characteristic (ROC) curves are provided in Sect. S.4.1.

### Analysis of TTs within 5° of myocyte minor axis

Because our method relies on convolution, quantification of local tubule orientation relative to the defined myocyte minor axis, termed tubule striation angle, is straightforward. Isolated cardiomyocytes and tissue sections were manually oriented via the included graphical user interface such that the observable majority of TTs were parallel to the y axis. We manually oriented the cardiomyocytes for simplicity, but in principle this process could be automated by utilizing angles inferred from the power spectral density of the low spatial frequency information of the outer SL. Filters were rotated at 5° increments, between −25° and 25°, and convolved with the image. TT filter hits at −5°, 0°, and 5° relative to the specified minor axis were compared to the total number of hits across all rotations.

### Tissue section analysis

Relative to analysis of isolated cardiomyocytes, the tissue-level characterization presented unique challenges, including segmention of non-myocyte regions. To address this challenge, we introduced a penalty filter to increase specificity of the TT filter in the presence of non-cardiac regions (see Sect. S.2.3 for more details). Thus, for these analyses, we did not mask the cell boundaries as done for the isolated cardiomyocytes. Given that no mask was applied to the images, we found that the performance of the LT filter and tubule striation angle analysis was substantially degraded and thus was not included.

### Computational tools

All aforementioned numerical procedures were conducted using the PYTHON2.7 libraries NUMPY, SCIPY and OPENCV-PYTHON. Data processing was performed using scipy and jupyter notebooks. All source code is provided at https://bitbucket.org/pkhlab/matchedmyo. Issue tracking and feature requests can be posted at https://bitbucket.org/pkhlab/matchedmyo/issues. A limited set of tests can be performed with the code at http://athena.as.uky.edu/node/7, however, for figure-quality images and custom analyses, download of the MatchedMyo software is strongly recommended. Details of the statistical analyses, algorithm use, installation, and algorithm refinements are provided in the supplemental methods. We have provided detailed installation and operation instructions in the Supplement (see Sect. S.1).

## RESULTS AND DISCUSSION

### Analysis of isolated cardiomyocytes

#### Cardiomyocytes from sham rat model

We first demonstrate the matched filtering approach using images of sham cells (Fig. 2A), in which the TTs exhibit a regular, striated pattern (see Sect. S.3.1 for details on cell isolation and imaging). Nevertheless, subregions of the cardiomyocyte presented an appreciable degree of spatial and angular variation. In the middle row of Fig. 2A, we present the classification results, for which the TT, LT, and TA responses are overlaid as blue, green, and red, respectively. In order to improve the classification results, we utilized a mask to exclude the cell membrane and nuclei in the field of view. In this image, the TT structure predominates over the LT and TA signals (middle row of Fig. 2A), which we summarize in Table 1. Finally, we present in the bottom row of Fig. 2A a heatmap of tubule striation angle superimposed onto the original image, where red corresponds to a TT filter hit at −25° and blue corresponds to a TT filter hit at +25°.

**Figure 2:**
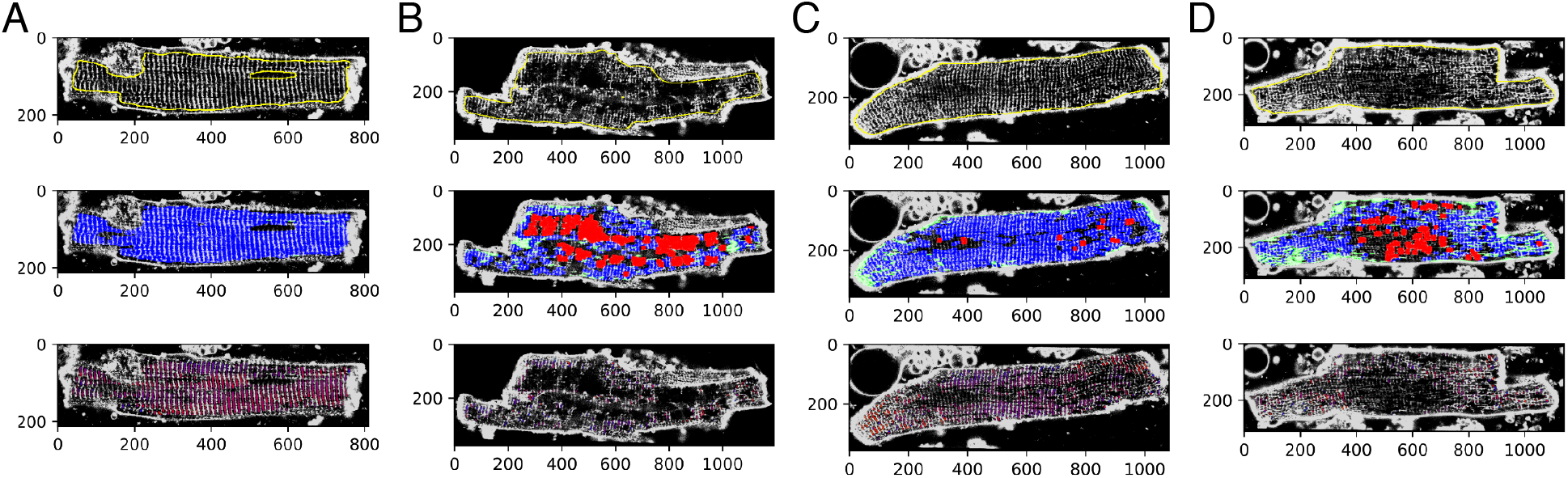
Algorithm application to isolated cardiomyocytes. Cardiomyocytes from (A) sham, (B) dilated cardiomyopathy via ascending aortic banding, (C) myocardial infarction (distal to infarct), and (D) myocardial infarction (proximal to infarct) models. (Top row) Preprocessed confocal images with outlines of image masks marked in yellow. (Middle row) MatchedMyo output with transverse tubule (TT), longitudinal tubule, and tubule absence filter hits marked in blue, green, and red, respectively. (Bottom row) Heatmap of tubule striation angle superimposed onto the image where red corresponds to TT hits at −25° and blue corresponds to TT hits at 25°.

**Table 1:** Quantification of content present within myocytes from differing disease etiologies, presented as percentage of cell area, *p < 0.05 compared to sham results of the same filter. TT denotes a “hit” from the transverse tubule filter. Similarly, LT and TA denote “hits” from the longitudinal and tubule absence filters, respectively. Distal and Proximal tissue results were analyzed using the bootstrapping method explained in Sect. S.3.4. Refer to the supplemental material for full range of cardiomyocytes considered herein. Data presented as mean ± SD.

| Case (n Cells) | TT (%) | LT (%) | TA (%) | TT $\pm 5^\circ$ Minor Axis (%) |
| --- | --- | --- | --- | --- |
| Sham (5) | $78 \pm 11$ | $4 \pm 2$ | $9 \pm 5$ | $57 \pm 12$ |
| AB (3) | $19 \pm 14^*$ | $5 \pm 1$ | $37 \pm 13$ | $36 \pm 9^*$ |
| MI<sub>Distal</sub> (3) | $56 \pm 15$ | $7 \pm 1$ | $11 \pm 6$ | $48 \pm 6$ |
| MI<sub>Intermediate</sub> (3) | $57 \pm 4^*$ | $10 \pm 2^*$ | $10 \pm 7$ | $37 \pm 4^*$ |
| MI<sub>Proximal</sub> (3) | $55 \pm 16$ | $12 \pm 3^*$ | $12 \pm 6$ | $34 \pm 4^*$ |
| Distal Tissue | $18.9 \pm 1.0$ | - | $9.5 \pm 2.0$ | - |
| Proximal Tissue | $10.1 \pm 0.8$ | - | $16.0 \pm 2.0$ | - |

In principle, similar conclusions can be drawn by visual inspection of these images or application of other analyses, such as the *TT_power_* method (see Sect. S.4.3 for a comparison of the MatchedMyo method to Fourier Transform TT density estimation). However, one benefit of the matched-filtering approach is the ability to detect spatially heterogeneous variations in the TT structure. For instance, the TT striations are not strictly straight throughout the cardiomyocyte interior. Rather, the striations appear to accommodate roughly perpendicular orientations relative to the outer SL along the major axis. In traditional TT index or power methods, such spatial variations in angular frequencies would present a diffuse signal response in the Fourier domain, leading to a potential misclassification of subcellular structure, unless outer sarcolemma is masked or the region of interest is restricted to a small area. Hence, our matched filtering approach is fundamentally different than techniques such as AutoTT[27], as our method computationally searches for arbitrary features within images, versus specifically characterizing transverse tubule periodicity or morphology. Further, because our method relies on convolution, we were able to quantify the angular variation (Table 1). Here, we find that sham cells present 57±12% of striations within 5° of the myocyte minor axis (see Sect. S.3.4 for details on statistical analysis). We also note that in rodent species, the T-system presents considerable branching that manifests in significant axial contributions [5]. These features were not evident in the cells considered here owing most likely to the lack of confocal z-stacks. For the LT and TA criterion, we report no significant responses in Fig. 2A.

#### Remodeling in AB cardiomyocytes

We next investigated T-system remodeling in tissue derived from a rat model of dilated cardiomyopathy based on AB [32]. In these animals, AB emulates pressure overload, which is known to drive hypertrophic remodeling broadly throughout cardiac tissue [33]. Commonly, tissue derived from AB mice present considerable TA, which correlates with impaired contractile function [28, 34]. Here we applied the algorithm to the tissue shown in Fig. 2B, using identical filters and thresholds from the previous sham analysis. As shown in Fig. 2B and Table 1, we find that TT content is significantly reduced (p<0.05) in AB isolated cardiomyocytes (19±14%) relative to sham (78±11%). This results in pervasive TA regions for AB cardiomyocytes (37±13%) vs. sham (9±5%). TA regions appear to be localized to the midline of the cell interior, whereas intact structure is mostly maintained adjacent to the outer plasma membrane. While TTs appear to be adjacent to the outer SL, there remains a significant (p<0.05) decrease in TTs within 5° of myocyte minor axis (36±9%) vs. sham. We report no significant changes in LT content for AB cardiomyocytes (5±1%) vs sham (4±2%).

#### Cardiomyocytes in MI rat heart

MI introduces a localized necrotic region in tissue. Cells adjacent to this necrotic region and border zone exhibit varying degrees of remodeling [35]. Here we applied the algorithm to cardiomyocytes isolated from regions distal, intermediate, and proximal to an MI, shown in Fig. 2C and D (medial cardiomyocytes not shown in figure, see Fig. S4 for all MI cardiomyocytes considered herein). These cells were obtained from an infarcted heart after explantation, for which the border zone was removed, and the remaining non-infarcted regions of the heart were divided into 3 equal-sized regions as described previously [36]. Relative to the sham cardiomyocytes, proximal cardiomyocytes exhibited a substantial increase in LTs (12±3%, p<0.05) as well as a decrease in TT density, indicated by TA filter hits (12±6%, Table 1). Cardiomyocytes from the intermediate region presented both LTs (10±2%, p<0.05) and TA (10±7%), although to a lesser extent than the proximal cells. Finally, we confirm for distally-derived tissue that for the four metrics used in this study, TT, LT, TA, and tubule striation angle, the differences observed were insignificant compared to sham cardiomyocytes. Relative to isolated cardiomyocytes from AB models, we see a substantial increase in LTs and a decreased amount of TA in MI. As with AB cardiomyocytes, a significant (p<0.05) decrease in the amount of TTs present within 5° of myocyte minor axis was observed in proximal tissue (34±4%) and intermediate tissue (37±4%) vs. sham. The main implication of this finding is that remodeling following MI appears to give rise to LTs and TT reorganization that is distinct from TA patterns exhibited in pressure overload models while exhibiting similar decreases in TTs within 5° of myocyte minor axis. Further, the extent of remodeling parallels distance from MI while TT density appears to have an inverse relationship with proximity to the MI.

### Analysis of cardiac tissue from infarcted rabbit heart

In Fig. 3, we present results for application of the MatchedMyo algorithm to MI tissue, which provides detail of the local tissue environment and the spatial distribution of tissue remodeling. Fig. 3A-C show portions of the infarcted tissue with no filtering, TT filtering, and TA filtering, respectively. Fig. 3B and C show heatmaps of TT and TA filter hits superimposed onto the tissue image. As observed previously in Fig. 2C and D, there was a marked decrease in the TT content, paralleled by an increase in TA detection, as the proximity to an infarct increased. Relative to the distal subsection, the proximal subsection (shown in further detail in Fig. S11C and D, respectively) presented, on average, 53.4% detection with the TT filter across the same area (see Table 1). This is paralleled by an increase in TA detection, with the proximal subsection presenting 68.4% more TA content than the distal subsection on average (see Sect. S.3.4 for further details on statistical analysis).

**Figure 3:**
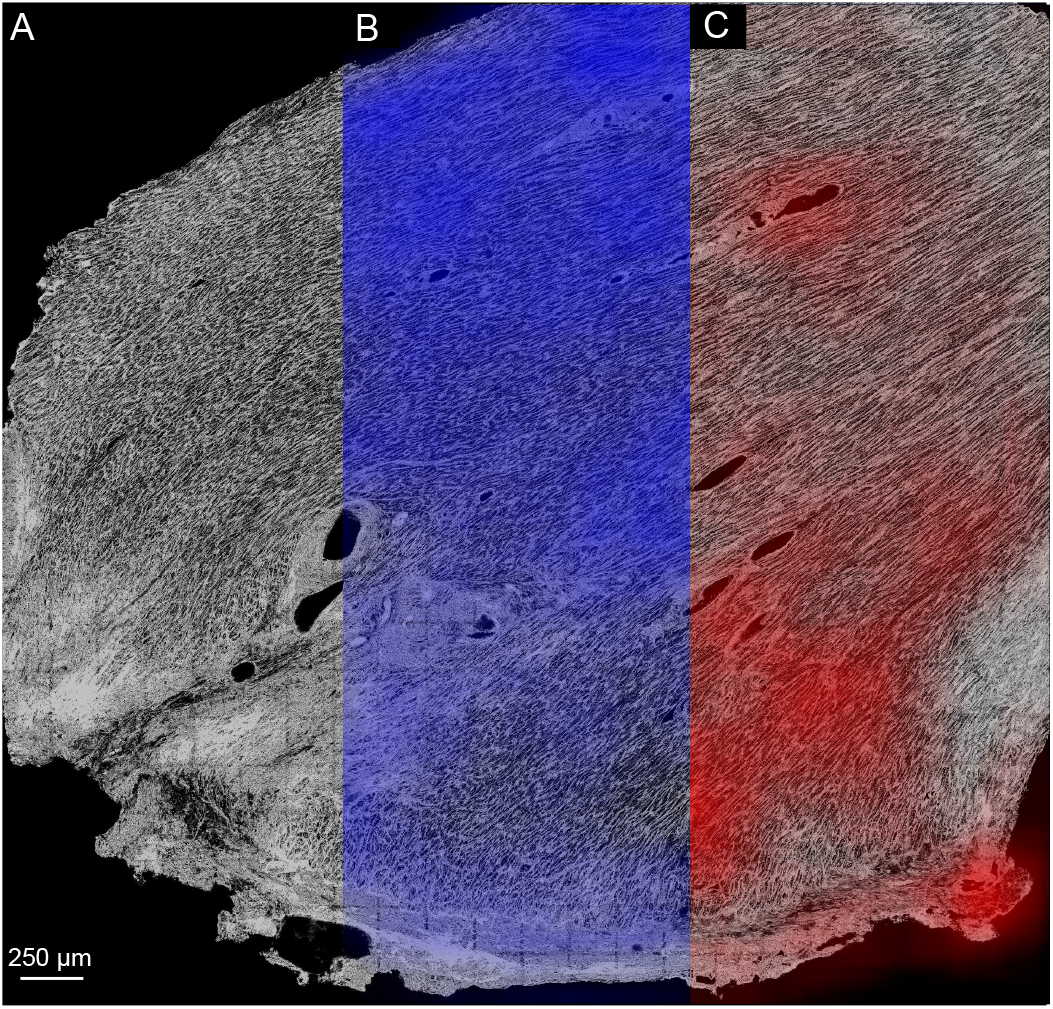
Application of TT filter (blue) and TA filter (red) to 3.9×4.1 mm^2^ sections of infarcted rabbit tissue where infarct is located at the bottom of the image, indicated by severe fibrosis. (A) Raw confocal tile scan exhibiting varying degrees of T-system remodeling. (B and C) TT and TA filter detections shown with heatmaps normalized to maximum density of filter detections. Striations arise from close apposition of punctate TT patterns typically observed in rabbit cardiomyocytes [37]. As summarized in Table 1, the distal region presented 18.9 =L· 1.0 % TT detection compared to 10.1 =L· 0.8 % in the proximal region. TA detection paralleled TT detection with the distal region exhibiting 9.5 =L· 2.0 % and the proximal region exhibiting 16.0 =L· 2.0 % TA detection.

## CONCLUSIONS

Our matched filtering-based technique complements existing computational algorithms for performing automated analyses of cardiomyocyte T-system [26, 27]. This approach leverages detailed structural information for the features of interest, based on kernels reflective of TT, LT, and TA morphology. Its reliance on convolution permits localized assessments of remodeling that scale to tissue-level preparations. We found that the approach performed remarkably well across cardiomyocytes from different species, disease etiologies, and fields of view. Our results suggest that the matched filtering approach performs favorably for detecting different extents and modalities for T-system configurations in confocal-microscopic images, and reveal patterns of T-system structure unique to sham, MI, and AB cardiomyocytes. We found that the approach is robust to regional variation in subcellular structure and noise, while providing a first-of-its-kind method scalable to tissue-level imaging.

For the cardiomyocytes we considered, the approach supported prior work revealing interesting trends in regional variation of cardiomyocyte T-system, such as the heterogeneity of TT organization in sham cells, as well as intracellular remodeling that depended on the distance from MI[36, 38]. We anticipate that these variations may give important clues into how cells adapt to their environment. This topic was previously explored by Guo et al [10], who cataloged diverse morphological configurations of cardiomyocyte T-system with respect to species and disease etiology.

Correlations between T-system disorganization and Ca^2+^ release dyssynchrony in numerous models of cardiac dysfunction [12, 35, 39, 40] suggest strong linkages between cardiomyocyte remodeling and pathophysiological function. It has been observed that SL proteins can redistribute under certain conditions that correlate with Ca^2+^ mismanagement, with no observation of increased TA[12]. Accordingly, it will be invaluable to investigate the potential correlations between T-system remodeling and protein localization, as well as the extent to which those relationships influence normal physiological function. In this regard, although the data considered in this study was limited to confocal images of dye-labeled SL, it is straightforward to adapt the algorithm to diverse data sets. Hence, in the event that filters representing features of interest can be identified, the approach can be adapted to a host of different labeling and imaging techniques, as well as cells isolated from a spectrum of tissue types.

## AUTHOR CONTRIBUTIONS

Following is a list of contributions made by each author. DFC: algorithm design, application/optimization of algorithm, wrote manuscript. SRB: algorithm design, edited manuscript. ACS: imaged tissue, edited manuscript. FBS: provided tools for tissue data acquisition, edited manuscript. MF: imaged isolated myocytes. WEL: provided tools for isolated myocyte data acquisition, edited manuscript. PKH: research design, algorithm design, wrote manuscript.

## ACKNOWLEDGEMENTS

This work was supported by the Maximizing Investigators’ Research Award (MIRA) (R35) from the National Institute of General Medical Sciences (NIGMS) of the National Institutes of Health (NIH) under grant number R35GM124977, as well as an Institutional Development Award (IDeA) from the National Institute of General Medical Sciences (NIGMS) of the National Institutes of Health (NIH) under grant number P20GM103527. DFC would also like to acknowledge support through the American Heart Association grant 17UFEL33490002. We thank Noah Adler for his support in integrating our utilities onto a web platform, Darin Vaughan and Ken Campbell for critical discussion of the manuscript, as well as Dr. Thomas McCoy for bootstrapping technique advice.

