## Supplement for "A Matched-filter Based Algorithm for Subcellular Classification of T-system in Cardiac Tissues"

### S.1 MATCHEDMYO INSTALLATION AND USAGE

The following sections lay out the installation and usage of the MatchedMyo software presented. To see an overview of the MatchedMyo classification algorithm visit `https://bitbucket.org/pkhlab/matchedmyo`.

#### S.1.1 Installation

Installation instructions are as follows.

**Unix-based work stations (Mac/Linux)**

Installation for Mac and Linux is identical. A basic knowledge of how to navigate using terminal is required.

1. *Mercurial installation.* Ensure that mercurial is installed on your work station. If mercurial is not installed, please click the link here for detailed installation instructions.

2. *Navigate to working directory/folder.* Open a terminal on your work station and navigate (using the '`cd`' command in terminal) to the directory/folder where you would like for the MatchedMyo software to be located.

3. *Pull the software from bitbucket to your work station.* Enter the command, "`hg pull https://bitbucket.org/pkhlab/matchedmyo`". At this point, the algorithm and some sample images will begin to download.

4. *Navigate to the new "matchedmyo" directory/folder.* Now, navigate to the new directory/folder created by the previous step.

5. *Download python libraries.* Choose to either automatically install python packages using PIP (preferred) or manually install the libraries. To automatically install the python libraries, type "`./installation.bash`" into the terminal and press "Enter". The libraries will automatically begin installing via PIP. If manual installation is wished, refer to the list of python libraries and versions in Sect. S.1.6.

6. *Generate filters.* Generate the filters needed for classification. To do this, execute the following command in terminal, "`python util.py -genAllMyo`". This generates all filters necessary for running the software and stores them in the "`myoimages`" folder of the `matchedmyo` folder.

7. *Verify installation.* The final step is to verify that the installation has completed without error. To do so, execute:
   "`python matchedmyo.py fullValidation`" The printing of "PASS" on the screen indicates that the MatchedMyo software has downloaded and installed correctly. If a "Runtime Error" is printed on the screen, the installation was not successful. This is likely due to library version inconsistencies. Run through the library installation instructions again and retry the validation. If this does not work, please email Dylan Colli at .

**Windows**

1. *Install the Anaconda Distribution for Windows.* The recommended way to install and use the MatchedMyo software suite is through installation of the Python 2.7 Anaconda Distribution. For the download link and installation instructions, visit `https://www.anaconda.com/download/#windows`.

2. *Open Anaconda Prompt.* In the Windows search bar, search for and launch the "Anaconda Prompt."

3. *Execute installation commands for Python libraries.* In the Anaconda Prompt, execute the following commands:
   pip install –user opencv-python=3.4.1.15
   pip install –user imutils==0.4.6
   pip install –user pygame==1.9.3
   pip install –user tifffile==2018.10.18

4. *Install Mercurial for Windows.* In the Anaconda Prompt, execute the following command:
   conda install -c conda-forge mercurial
   Confirm the packages to be installed by typing 'y' and pressing 'Enter.'

5. *Navigate to folder that will house algorithm.* Change your working folder in the Anaconda Prompt by using the 'cd' command. It is recommended that you change your working folder to your desktop or something equally as accessible. An example of this command execution is as follows:
cd C:\Users \Dylan \Desktop
However, individual paths to the desktop will vary based on username.

6. *Pull the software from bitbucket to your work station.* To download the repository from bitbucket, execute the following command in the Anaconda Prompt:
"`hg clone https://bitbucket.org/pkhlab/matchedmyo`"

7. *Change your working folder to the MatchedMyo folder.* To change your working folder, execute, `cd matchedmyo`.

8. *Generate filters.* Generate the filters needed for classification. To do this, execute the following command: "`python util.py -genAllMyo`"

9. *Verify installation.* The final step is to verify that the installation has completed without error. To do so, execute:
"`python matchedmyo.py fullValidation`" The printing of "PASS" on the screen indicates that the MatchedMyo software has downloaded and installed correctly. If a "Runtime Error" is printed on the screen, the installation was not successful. This is likely due to library version inconsistencies. Run through the library installation instructions again and retry the validation. If this does not work, please email Dylan Colli at .

#### S.1.2 Command line usage for TT, LT, and TA filters

MatchedMyo is ran via the command line. To do this, follow these steps:

1. *Open a terminal.* MatchedMyo is operated through the command line interface. To access the command line interface on 1) Linux distributions, open the Dash and type "Terminal" into the search bar, 2) Mac, open Spotlight and search for "Terminal", and 3) on Windows, launch the Anaconda Prompt.

2. *Navigate to the directory/folder containing the MatchedMyo software.* Next, we must navigate to the directory or folder containing the MatchedMyo software. Change directories in terminal or the Anaconda Prompt using the 'cd' command.

3. *Create/edit a YAML (.yml) file.* Non-default parameters are specified through YAML files when running MatchedMyo. One must either create a fresh YAML file or edit the template.yml file included in the "`YAML_files`" folder of the MatchedMyo software. Only one parameter, the image name, is required. To specify this, include "`imageName: <NAME_OF_IMAGE>`" in the YAML file, where "`<NAME_OF_IMAGE>`" is the path to, and the name of, your image. To specify other parameters, include "`<PARAMETER_NAME>: <PARAMETER>`" in the YAML file, where "`<PARAMETER_NAME>`" is the name of the parameter and "`<PARAMETER>`" is the value of the parameter you would like to specify. For a full list of parameter options and acceptable values, see Sect. S.1.4.

4. *Run the MatchedMyo file.* This is accomplished by executing the following command in terminal, "`python matchedmyo.py run --yamlFile <YAML_FILE>`". Where "`<YAML_FILE>`" is the name of the YAML file generated in the previous step.

5. *View outputs.* If `fileRoot` is specified in the YAML file, the classified images will be output to that location. Additionally, the analysis of TT, LT, and TA content will be stored in a comma-separated values (CSV) file. The default location for this CSV file is under the "results" folder in the MatchedMyo software folder, but a new location can be specified using the YAML file. See Sect. S.1.4 for full instructions on how to specify parameters via the YAML file.

#### S.1.3 Command line usage for arbitrary filters

As explained in the README file of the repository hosted on bitbucket, the algorithm may be applied to arbitrary images and filters using the script MATCHEDMYO.PY. Now, instead of relying on default parameters such as in the case for TT, LT, and TA filtering, the parameter dictionaries used for classification (`paramDicts` in Sect. S.1.4) must now be fully specified in the YAML file. The parameter dictionaries' structures are unmodified except for the name that is used to specify them. Instead of using 'TT,' 'LT,' and 'TA,' one now uses 'filter1,' 'filter2,' and 'filter3' where 'filter1' hits are marked in blue, 'filter2' hits are marked in green, and 'filter3' hits are marked in red. Additionally, one must add "`classificationType: arbitrary`" to the YAML file. The following is an example YAML file.

```
imageName: myoimages/Sham_11_processed.png
classificationType: arbitrary
outputParams:
  fileRoot: filterHits
paramDicts:
  filter1:
    filterMode: punishFilter
    filterName: ./myoimages/newSimpleWTFilter.png
    punishFilterName: ./myoimages/newSimpleWTPunishmentFilter.png
    snrThresh: 0.35
    gamma: 3
```

We provide an example output in Fig. S1.

#### S.1.4 List of YAML parameters for MatchedMyo classification

The following is a list of all tunable parameters/options available through the YAML file functionality.

**User parameters** - The following are parameters used to tune what classification is performed and what is output by the software.

- imageName - Path to, and name of, the image that is to be classified by the MatchedMyo software.
  - *Acceptable Inputs* - Any string.
  - *Example Input* - imageName: /home/user1/Pictures/image1.png
- outputParams - This is how the output file name, file type, dots per inch (DPI, resolution) of the output images, and the classification results CSV file is specified. If the file name is None, like the default input, then the output files are not written. The default storage location for the classification results CSV file is located in the "results" folder of the MatchedMyo software. This CSV file contains information such as date and time of analysis, name of the image, TT, LT, and TA content, and output image location and name.
  - *Acceptable Inputs* -
    ```
    outputParams:
        fileRoot: <a string>
        fileType: <png, tif, or pdf>
        dpi: <an integer>
        saveHitsArray: <True or False>
        csvFile: <a string>
    ```
  - *Default Input* -
    ```
    outputParams:
        fileRoot: None
        fileType: png
        dpi: 300
        saveHitsArray: False
        csvFile: ./results/classification_results.csv
    ```
- preprocess - This turns preprocessing on or off.
  - *Acceptable Inputs* - preprocess: <True or False>
  - *Default Input* - preprocess: True
- filterTypes
  - *Acceptable Inputs* -
    ```
    filterTypes:
        TT: <True or False>
        LT: <True or False>
        TA: <True or False>
    ```
  - *Default Input* -
    ```
    filterTypes:
        TT: True
        LT: True
        TA: True
    ```

- `scopeResolutions` - This is the resolution of the confocal microscope used to collect images in pixels/voxels per micron. Note that this uses the convention of "x" being the first axis, corresponding to the rows of the image. Thus, "y" corresponds to the columns in the image.
  - *Acceptable Inputs* -
    ```
    scopeResolutions:
        x:  <an integer or float>
        y:  <an integer or float>
        z:  <an integer, float, or None>
    ```
  - *Default Input* - None
  - *Example Input* -
    ```
    scopeResolutions:
        x:  5.03
        y:  5.03
    ```
- `iters` - This parameter controls the rotations at which the filters will be analyzed.
  - *Acceptable Inputs* - `iters:  <a list of integers>`
  - *Default Input* - `iters:  [-25, -20, -15, -10, -5, 0, 5, 10, 15, 20, 25]`
- `returnAngles` - This option turns on/off the analysis and output of transverse tubule striation angle.
  - *Acceptable Inputs* - `returnAngles:  <True or False>`
  - *Default Input* - `returnAngles:  False`
- `returnPastedFilter` - This option turns on/off the superimposing of filter-sized unit cells on the hits of the original image. If this is turned off, only the original 'hits' will be superimposed on the image. If this is turned on, for each hit, a filter-sized rectangle will be placed on the image to mark the filter on the image. Keeping this option on is much more intuitive.
  - *Acceptable Inputs* - `returnPastedFilter:  <True or False>`
  - *Default Input* - `returnPastedFilter:  True`

**Developer parameters** - The following is a list of parameters that the ordinary user does not need to change. However, if a user wishes to change filters, detection strategies, or filter thresholds, this is how it is done.

- `filterTwoSarcomereSize` - This parameter designates the filter size in relation to the myocytes/tissue we wish to analyze. Unless experimenting with non-default filters, this should be left alone.
  - *Acceptable Inputs* - `filterTwoSarcomereSize:  <an integer>`
  - *Default Input* - `filterTwoSarcomereSize:  25`
- `paramDicts` - This holds all of the parameters needed for each individual filtering routine (TT, LT, and TA). `filterMode` refers to the type of SNR calculation being utilized for the classification. There are three options for this parameter, `simple` just calculates the convolution of the filter with the image, `regionalDeviation` utilizes the SNR calculation explained in Sect. S.2.3, "Penalty against filter with spectral overlap," and `punishmentFilter` utilizes the SNR calculation explained in Sect. S.2.3, "Standard deviation criterion." `filterName` is the name of the filter used for each morphological classification. `punishFilterName` is the name of the 'punishment' filter used if `filterMode:  punishmentFilter` is specified. `gamma` is the scalar used to weight the punishment filter response. `snrThresh` is the threshold used to differentiate classification "hits" versus "non-hits" using the signal to noise ratio. `stdDevThresh` is the threshold used differentiate classification "hits" versus "non-hits" using the standard deviation criterion used in the `regionalDeviation` filter mode. `inverseSNR` refers to the flag that indicates whether the classification "hits" are above the `snrThresh` or below. False indicates that classification "hits" are above the threshold, True indicates "hits" are below the threshold.
  - *Acceptable Inputs* -
    ```
    paramDicts:
    ```

```
TT:
     filterMode:  <simple, regionalDeviation, punishmentFilter>
     filterName:  <a string>
     punishFilterName:  <a string>
     gamma:  <an integer or float>
     snrThresh:  <an integer or float>
LT:
     filterMode:  <simple, regionalDeviation, punishmentFilter>
     filterName:  <a string>
     snrThresh:  <an integer or float>
     stdDevThresh:  <an integer or float>
TA:
     filterMode:  <simple, regionalDeviation, punishmentFilter>
     filterName:  <a string>
     inverseSNR: <True or False>
     snrThresh:  <an integer or float>
     stdDevThresh:  <an integer or float>
```

– *Default Input* -

```
paramDicts:
   TT:
        filterMode:  punishmentFilter
        filterName:  ./myoimages/newSimpleWTFilter.png
        punishFilterName:  ./myoimages/newSimpleWTPunishmentFilter.png
        gamma:  3.0
        snrThresh:  0.35
   LT:
        filterMode:  regionalDeviation
        filterName:  ./myoimages/LongitudinalFilter.png
        snrThresh:  0.6
        stdDevThresh:  0.2
   TA:
        filterMode:  regionalDeviation
        filterName:  ./myoimages/LossFilter.png
        inverseSNR: True
        snrThresh:  0.04
        stdDevThresh:  0.1
```

##### S.1.5 Webserver

As stated in the Methods, we freely provide access to our python implementation of this approach at http://athena.as.uky.edu/node/7. The online implementation permits uploading of trial image files for testing the most basic functions of the MatchedMyo software. It is important to note, however, that this is a simplified version of the MatchedMyo software and if figure-quality images and analysis are desired, installation of the MatchedMyo software on a local work station is strongly recommended. It is also worthy to note that although little pre-processing was done in our study, user-provided data may need some corrections to ensure consistent illumination and noise reduction, as well as compatible filter and data spatial scales. More advanced analysis, including retraining of the detection data, will require download of the bitbucket repository.

**Usage** - Upon visiting http://athena.as.uky.edu/node/7, the user will be prompted to upload the image file to be classified, an optional mask, an optional YAML file and will be asked to enter an email address where a download link for the classification results will be sent. For information on creating an image mask, see Sect. S.3.2. The YAML file follows the same conventions and can be modified in the same manner as outlined in Sect. S.1.4, except for the `preprocess` parameter option. For the webserver implementation, if `preprocess` is specified as "True" (such as in the default case), the image will go through a series of automatic preprocessing routines. These routines are manually performed by the user when running the MatchedMyo software locally, however this implementation is not possible through the webserver interface. This automatic preprocessing is less tolerant of noise, but is convenient for rapid prototyping of MatchedMyo application through the webserver. Aside from this parameter, the default webserver YAML parameters are those used to produce the figures herein.

Upon submitting the job request, the user will receive an email with a link to a color-coded image indicating the filter responses. The email should be expected within one to two minutes of submitting the job.

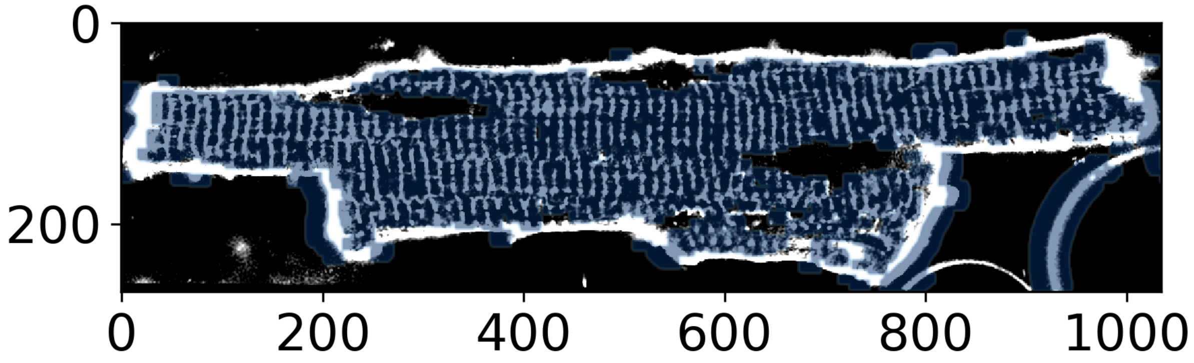


Figure S1: Cardiomyocyte (black and white) for which filter responses from the arbitrary filtering scheme are marked in blue.

#### S.1.6 List of Python libraries and version numbers

The following is a list of Python libraries used in the MatchedMyo software suite.

- OpenCV
  - PIP library name: opencv-python
  - Version number: 3.4.1.15
- Imutils (https://github.com/jrosebr1/imutils)
  - PIP library name: imutils
  - Version number: 0.4.6
- Numpy
  - PIP library name: numpy
  - Version number: 1.15.4
- Matplotlib
  - PIP library name: matplotlib
  - Version number: 2.2.3
- Tkinter (default Matplotlib Pyplot GUI backend)
  - Library name: python-tk
  - Version number: 8.6

- PyYAML
  - PIP library name: PyYAML
  - Version number: 3.12
- Scipy
  - PIP library name: scipy
  - Version number: 1.1.10
- Pandas
  - PIP library name: pandas
  - Version number: 0.23.4
- Pygame (https://www.pygame.org/download.shtml)
  - PIP library name: pygame
  - Version number: 1.9.3
- Python Image Library (PIL)
  - PIP library name: Pillow
  - Version number: 5.2.0
- SciKit Learn
  - PIP library name: scikit-learn
  - Version number: 0.19.1
- Tifffile
  - PIP library name: tifffile
  - Version number: 2018.10.18

### S.2 SUPPLEMENTARY THEORY

#### S.2.1 Detection by convolution

Beyond classifying whether cellular microstructure resembles known filters, our workflow is tasked with designating regions of T-system absence as well as unclassified TT structure. For this reason, we present the following detection scheme. We assess the likelihood of subcellular content via

$$\bar{y}_i = \hat{h}_i * \hat{m} > \lambda_i, \quad \forall\, \hat{h}_i \; in \; \{\hat{h}_{TT}, \hat{h}_{LT}\} \tag{S1}$$

$$\bar{y}_{TA} = \hat{h}_{TA} * \hat{m} < \lambda_{TA} \tag{S2}$$

where $*$ represents the convolution of the filter, $\hat{h}_{TT}$, $\hat{h}_{LT}$, and $\hat{h}_{TA}$, with the confocal microscopy image, $\hat{m}$, and $\lambda_i$ represents the user-determined threshold. A region is considered uncharacterized if no $\bar{y}_i$ is indicated as a hit via this criterion.

#### S.2.2 Filter bank

In our approach, we additionally consider filter rotations to detect alternate tubule orientations through defining a bank of filter rotations for each matched filter spaced at 5 degree increments. The optimal rotation for a given filter is determined by evaluating each rotated filter with the data; the rotated filter that generates the highest above-threshold response is elected as the most likely orientation of a given feature, $j$:

$$y_i = \max(y_{i,\phi_j}), \quad \forall\, \phi_j \in \{\phi_0, \phi_1, \ldots, \phi_n\} \tag{S3}$$

$$y_{i,\phi_j} = (R(\phi_j)h)^T m \tag{S4}$$

where $R(\phi_j)$ represents the rotation of filter $h$ by $\phi_j$.

#### S.2.3 Additional refinements

**Penalty against filters with spectral overlap** Since there is considerable similarity between the longitudinal and transverse tubule matched filters used in this approach, we propose a penalty term to reduce false positives arising from spectral overlap of the intended signal with an undesired filter (e.g. TT versus LT). Hence, we modify Eq. 1 to include a penalty term, $p$, based on the power spectral density of undesired filters. This yields the revised equations

$$\mathbf{C_n}' = \mathbf{C_n} + p\mathbb{E}\{\hat{h}_u \hat{h}_u^T\} \tag{S5}$$

$$\hat{h} = \frac{1}{(\hat{s}^T \mathbf{C_n}'^{-1} \hat{s})^{\frac{1}{2}}} \mathbf{C_n}'^{-1} \hat{s}, \tag{S6}$$

where $\hat{h}_u$ represents the undesired filter. This complement penalizes signal that falls outside of the signal signature defined in $\hat{h}_i$. We tested both filters against annotated data in Sect. S.4.1 and demonstrate this formulation is necessary to discriminate the correlation outputs from the longitudinal and TT regions.

**Standard deviation criterion** To further increase specificity for both LT and TA filters, we found it necessary to include a standard deviation criterion. That is, for a region to be considered a hit with the longitudinal filter, the region must satisfy 1) super-threshold response of the general matched filtering protocol described previously, and 2) sub-threshold response of a convolution standard deviation measurement. The second criterion was calculated as previously described,[1] the summary of which is as follows.

$$\sigma = (\frac{1}{N-1}(q - \frac{s^2}{N}))^{\frac{1}{2}} \tag{S7}$$

$$q = \sum_{i=1}^{N} x_i^2 \tag{S8}$$

$$s = \sum_{i=1}^{N} x_i, \tag{S9}$$

where $\sigma$ represents the standard deviation of the data, $x$ is the complete data set, $x_i$ represents the region of interest in $x$, and $N$ is the number of data points within $x$. Since $q$ and $s$ are summations, they can be calculated by convolving $x^2$ and $x$, respectively, with a kernel of 1's with equal dimensions as the region of interest (the area of the longitudinal or TA filter for this case).

#### S.2.4 Matched filter training and testing

The filters used in the paper were idealized representations of the signals we anticipated to observe in the longitudinal and transverse tubule morphologies with minor alterations in the longitudinal filter to account for spectral overlap between the two tubule filters. That said, we have also performed tests using filters that are directly determined from the data, by averaging the transverse or longitudinal features we identified by visual inspection in the isolated myocytes. While we found that the idealized filters performed more robustly, the matched filtering approach can use either ad hoc or data-informed filters. We will also elaborate on a distinction between our implementation of the matched filtering approach and the basic theory of matched filtering. As suggested in the Wiki reference, it can be shown that the matched filter maximizes the SNR when presented with measured data (m) comprised of a signal (s) and additive, uncorrelated noise (n), e.g. m = s + n . In practice, the signal (transverse tubules) is not strictly homogeneous and the noise can have significant nonlinearity and correlation; nevertheless, and as our data indicate (particularly the sensitivity analyses/ROC curves), the matched filtering approach can yield significant and accurate detections for filters/signals that reasonably approximate the feature one intends to identify.

After the filters were determined, two sets of parameters were optimized: 1) the contribution of the penalty filter weight, $p$, in Eq. S5, 2) the standard deviation threshold described in Sect. S.2.3, and 3) the detection thresholds, $\lambda_i$, in Eq. S1 and Eq. S2. Details for selecting optimal threshold parameters and corresponding receiver operator characteristic (ROC) curves are provided in Sect. S.4.1. Namely, the optimization consisted of evaluating algorithm performance using ROC curves, for which the model parameters were varied at regular intervals across a range of values. Optimal parameters were chosen such that the normalized true positive rate was maximized while the normalized false positive rate was constrained to below 0.25 (see Table S3 for values).

### S.3 SUPPLEMENTARY METHODS

Details for the confocal image acquisition and algorithm workflow are elaborated below. Algorithm workflow and its implementation as a webserver is provided in Sect. S.1.5.

#### S.3.1 Isolated cell and tissue preparation

Di-8-ANEPPS-labelled isolated cells were imaged, segmented, and deconvolved (to suppress noise) according to standard procedure (as described in [2–7]). In this study, we consider isolated cardiomyocytes from two disease Wistar rat models, dilated heart failure with reduced ejection fraction induced via AB (Fig. S5), and MI (Fig. S4)[3]. We additionally examine whole-tissue preparations from an infarcted rabbit (see Fig. 3). Rabbit infarct tissue was excised and labeled with CF488A-conjugated wheat germ agglutinin as described previously [8]. These data reveal a preponderance of linear striations evident of the t-system. For diseased models, this structure is either lost or remodeled with longitudinal tubules [9]. Since the primary goal of the matched filtering is to discriminate between different types of remodeling, the underlying physiological reasons for their effects on calcium homeostasis are not considered.

#### S.3.2 Input

Input consists of 1) either a unicellular or multicellular fluorescence confocal microscopy image and 2) a user generated mask if the image is unicellular. Both images can be either the Portable Network Graphics (png) or the Tagged Image File Format (tif) file types. The mask convention is such that all intracellular pixels are marked as the image's brightest pixel intensity and all non-intracellular pixels are less than the brightest pixel intensity. The mask is to be generated in the user's choice of image editor. At this point, the user initiates the algorithm from the command prompt, designating the paths to both the confocal data and mask.

#### S.3.3 Preprocessing

**Image reorientation** The first step of the algorithm consists of the user drawing a line orthogonal to the transverse tubule network using the supplied graphical user interface (GUI). This line serves as a basis for re-orientation of the major axis of the myocyte(s). This manual reorientation allows for explicit reorientation as opposed to automatic methods of reorientation such as principal component analysis that are sensitive to extracellular content and thus may misalign if the images are not carefully segmented. Additionally, reorienting the myocyte(s) limits the bank of filter rotations the algorithm must consider, considerably reducing computational expense. However, this reorientation is not a strict requirement to obtain high accuracy detections, as we demonstrate with multi-cellular preparations in Fig. 3.

**Image resizing** Secondly, the user may select a subsection of the image that contains as many conserved striations as possible without including membrane or highly heterogeneous regions. From this subsection, a frequency analysis is calculated to determine the length (measured in pixels) per striation. This value is used to resize the image to a common length scale. In the event that the spatial resolution of the image is known and from which the sarcomere length can be estimated, this information can be used directly to rescale the image.

**Application of Contrast-Limited Adaptive Histogram Equalization (CLAHE)** Further, the previously discussed histogram equalization technique is implemented via OpenCV's CLAHE routine [10] to normalize variations in signal intensity that we attributed to uneven immunofluorescence labeling.

**Normalization of fluorescent intensity to tubules** Finally, TT normalization was implemented to obtain consistent classification across datasets that may contain fluctuations of TT fluorescence between images. This routine utilizes OpenCV's Gaussian thresholding to obtain a rough approximation of TT striation and intracellular fluorescent intensity. Each image was thresholded and normalized according to these values such that all pixels above the estimated TT maximum are marked as the maximum and all pixels below the estimated minimum are marked as the minimum. The image is then normalized for algorithm use.

#### S.3.4 Statistical analysis

Statistical data (mean content and standard deviation) presented in the main text for isolated myocytes are based on 5 cell images for the sham case and 3 cell images for the AB and each proximity of the MI case. Figures provided are based on a representative member of each set of $n$ images that are displayed in Fig. S3 (sham), Fig. S4 (MI), and Fig. S5 (AB). For the multi-cellular tissue preparation, we utilized a bootstrapping method for statistical analysis as only one raw image was available. The image was divided into three sections based on proximity to the infarct. The upper one-third of the tissue-level image was designated distal to the infarct, the middle one-third intermediate, and the lower one-third, closest to the infarct, was deemed proximal to the infarct. After applying an exclusion criterion such that any 2000x2000 subimage that comprised less than three-quarters cardiomyocytes was discarded, the distal region comprised 22 subimages and the proximal region comprised 18 subimages. The distal subimage population was pared down to match the population size of the proximal population, 18 subimages, randomly using numpy's random module. From these subimages, we constructed 1000 bootstrap images consisting of selections from 25 randomized 400x400 subsamples of the 18 original 2000x2000 (.35x.35 mm$^2$) subimages for the respective proximal and distal regions. The random selections were based on a draw and replacement [11] method, whereby a given subsample could be selected more than once. Each of the 1000 bootstrap images were then analyzed using the matched filtering algorithm, from which mean TT and TA content were reported in Table 1.

Table S1:

| Proximity to Infarct | TT Detection Rate (%) | TA Detection Rate (%) |
|---|---|---|
| Distal | 18.9 ± 1.0 | 9.5±2.0 |
| Proximal | 10.1 ± 0.8 | 16.0±2.0 |

To determine statistical significance, we assessed the difference of means for each bootstrap image,

$$\Delta\bar{X}_i \equiv \bar{X}_{i,1} - \bar{X}_{i,2}, \quad \text{for } i = 0, \ldots, 1000 \tag{S10}$$

From these means, the confidence interval for $\Delta\bar{X}$, was obtained by rank ordering the variates $\{\Delta\bar{X}_i, ..., \Delta\bar{X}_n\}$, with $\bar{X}_i \leq \bar{X}_i \leq \cdots \leq \bar{X}_n$. The 95% confidence interval is defined by the 2.5th and 97.5th percentiles variates, e.g.

$$[\Delta\bar{X}_{i=25}, \Delta\bar{X}_{i=975}]$$

for N=1000 variates. If this interval does not contain $\Delta\bar{X} = 0$, then the means are assumed to be statistically different with 95% certainty. The confidence interval for TT and TA content is reported in Table S2

Table S2: Results for difference of means calculation applied to rabbit infarct model for transverse tubule (TT) and tubule absence (TA) criterion. Range of variates calculated for p<0.05. The absence of zero within $[\Delta\bar{X}_{i,25}, \Delta\bar{X}_{i,975}]$ indicates 95% confidence that criterion i possesses dissimilar means for the proximal and distal regions of the infarcted tissue.

| | $[\Delta\bar{X}_{TT,25}, \Delta\bar{X}_{TT,975}]$ (%) | $[\Delta\bar{X}_{TA,25}, \Delta\bar{X}_{TA,975}]$ (%) |
|---|---|---|
| Difference of Means | $[6.1, 12.0]$ | $[-12.0, 0.0]$ |

The observed magnitude in the difference of means for measured TT and TA content is diminished for bootstrapped images compared to isolated cardiomyocyte images due to several factors. For the tissue-level image, non-myocyte-containing regions are introduced and overrepresented during the bootstrap image construction and analysis. Blood vessels and significant fibrosis, the former of which was prominent in the distal region, simultaneously attenuated the response of the TT filter and increased the response of the TA filter. Additionally, image artifacts in the form of edge-effects are introduced in the stitching of bootstrap images. Filters are truncated along the borders of images which introduces numerous false positives for the TA filter. Stronger performance and statistical significance would be expected to arise from the use of multiple tissue-level infarct images.

### S.4 SUPPLEMENTARY RESULTS

#### S.4.1 Matched filter detection protocol and performance

For the detection criterion in Eq. S1 and Eq. S2, we defined 'true positive' and 'false positive' metrics that we sought to maximize or minimize, respectively, for a given choice of ($\lambda_{\text{TT}}$, $\lambda_{\text{LT}}$, and $\lambda_{\text{TA}}$). These metrics were evaluated for the three phenotypes considered in this study a) sham, b) MI and c) AB. Each image was manually annotated to indicate possible regions with characterized tubule structure (TT and LT), or tubule absence (TA). The true positive metric was based on enumerating above-threshold pixels for a given filter that overlapped with the hand-annotated markers. The false positive metric enumerated above-threshold pixels that appeared in regions not marked for a given filter. Because the manual annotations were imprecise and the false positive pixels frequently scaled with image size, we normalized the true positive and false positive rates by the pixel counts returned for the lowest threshold parameters considered. Hence, for $\lambda_{\text{TT}} = 0$ the comparison would return normalized true positive and false values of 1.0, though the unnormalized values would typically be much larger for the true positive relative to false positive pixels. In Fig. S2 we present ROC curves to assess the relative true positives (correct detections) versus false positive (incorrect detections) as a function of a cutoff parameter, $\lambda$. The choice of $\lambda_i$ in $\lambda_a = 0.2$ is intended to balance high true positive rates while minimizing false positives, and is in principle an application-specific parameter. For our data, we selected $\lambda_{WT} = 0.35$, $\lambda_{LT} = 0.6$, $\lambda_{TA} = 0.04$ for the three filters. The true positive and false positive rates for each of the annotated images and filters is displayed in Table S3. The data considered in this study was reasonably high contrast, thus we would expect the performance to degrade as a function of noise amplitude. We note that simulated data could be used to optimize detection criteria, as was done in [12], for instance. We opted rather to use real imaging data, as there are numerous sources of variation that are difficult to reproduce via simulation. Among these include random variations in the TT morphology (diameter, length, orientation), heterogeneities in fluorescent indicator distribution, and noise characteristics specific to the sensor that may not necessarily be non-additive, as assumed in Eq. 2. For this reason, optimization to simulated data is likely to yield criteria that would not perform well with real data.

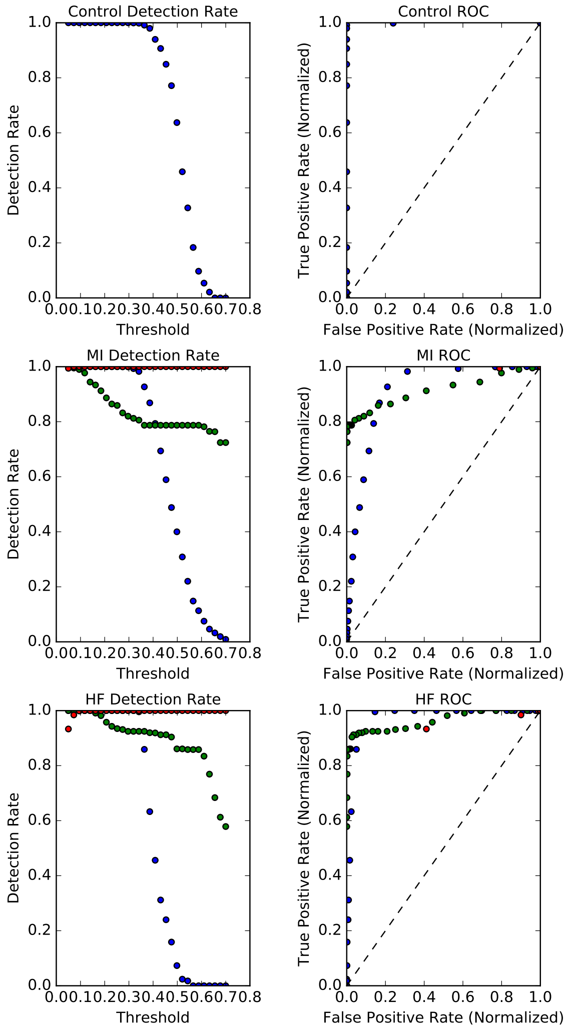


Figure S2: True positive rates (left) and ROC curves (right) for hand annotated control/sham data (top), infarct data (middle), and ascending aortic banding data (bottom). Blue points represent the transverse tubule filter, green represent the longitudinal tubule filter, and red represent the tubule absence filter. Absence of longitudinal tubule and tubule absence data in the control ROC curve is due to the lack of those morphological features in the training data. Filters present high performance across the three training sets as shown by the high area under each ROC curve.

Table S3: True positive and false positive rates for native transverse tubule structure (TT), longitudinal tubule (LT), and tubule absence (TA) filters. - indicates that no feature of interest for that particular filter was present in the annotated image.

| Annotated Image | $WT_{TP}/WT_{FP}$ | $LT_{TP}/LT_{FP}$ | $TA_{TP}/TA_{FP}$ |
|---|---|---|---|
| Sham | 0.996/0.070 | -/- | -/- |
| MI | 0.970/0.249 | 0.695/0.002 | 0.909/0.002 |
| AB | 0.973/0.107 | 0.806/0.002 | 0.859/0.0001 |

#### S.4.2 Analysis of Three Dimensional Simulated Cells

Table S4: Quantification of content present within myocytes from differing disease etiologies, presented as percentage of cell area. TT denotes a "hit" from the transverse tubule filter. Similarly, LT and TA denote "hits" from the longitudinal and tubule absence filters, respectively. Note that Fig. S7 results were normalized to the conserved striation tissue subsection. Data presented as mean ± SD.

| Figure | Case | TT (%) | LT (%) | TA (%) | TT ± 5° Minor Axis (%) |
|---|---|---|---|---|---|
| Fig. 2A | Sham | 95 | 0.2 | 1 | 65 |
| Fig. 2B | AB | 37 | 6 | 29 | 38 |
| Fig. 2C | $MI_{Distal}$ | 77 | 7 | 3 | 55 |
| Fig. 2D | $MI_{Proximal}$ | 39 | 16 | 14 | 32 |
| Fig. S7 | Distal Tissue | 100 | - | 100 | - |
| | Proximal Tissue | 45.8 | - | 574 | - |

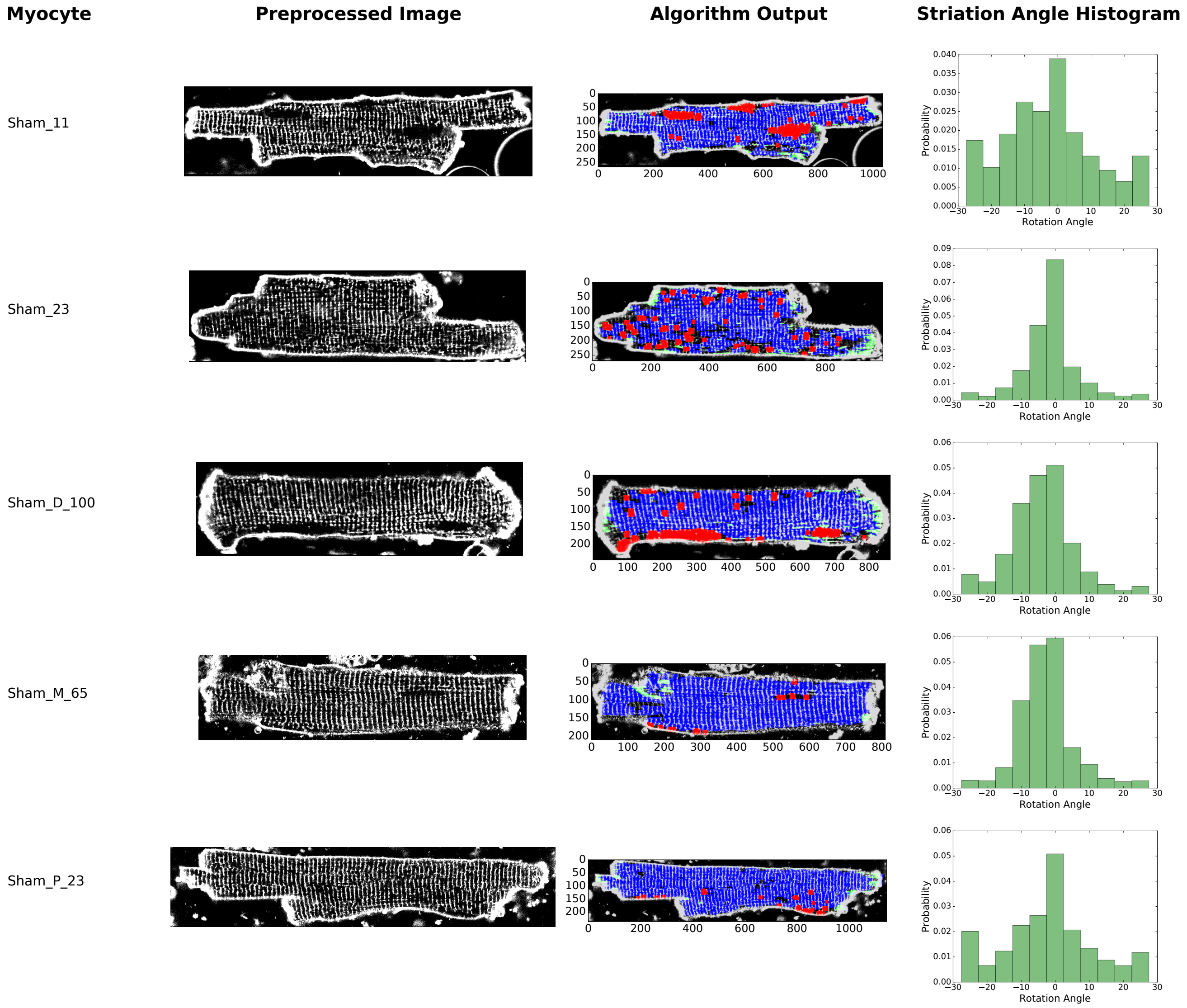


Figure S3: Results for all isolated sham myocytes considered in this study. Isolated myocyte file name (leftmost column). Preprocessed image (second column from left). Algorithm output marking transverse tubules, longitudinal tubules, and tubule absence in blue, green, and red, respectively (second column from right). Histogram of transverse tubule detection angle (rightmost column).

A strength of the matched filtering method is the technical ease of extension from classification in two dimensions to three dimensions. Incorporation of another dimension of the confocal image, termed z-stacks, leads to a more robust calculation due to integration of another dimension of measured signal. Thus, as a proof-of-concept for three dimensional classification we have included analysis of simulated cells in Fig. S6 and Table S8. Simulated cells were constructed based upon sequential addition of unit cells for TT, LT, and TA morphologies given user-defined probabilities of finding each morphological feature. Twenty simulated cells were generated and analyzed

Table S5: Quantification of content present within sham myocytes, * indicates the myocyte was analyzed with a mask for a region of interest excluding an assumed organelle. TT denotes a "hit" from the transverse tubule filter. Similarly, LT and TA denote "hits" from the longitudinal and tubule absence filters, respectively.

| Myocyte | TT (%) | LT (%) | TA (%) | % TT Hits Within 5° of Minor Axis |
|---|---|---|---|---|
| Sham_11 | 73.0 | 5.1 | 13.1 | 40.9 |
| Sham_11* | 83.8 | 2.1 | 7.3 | 42.4 |
| Sham_23 | 62.9 | 6.3 | 12.4 | 73.0 |
| Sham_D_100 | 73.0 | 5.4 | 12.3 | 57.7 |
| Sham_D_100* | 83.3 | 2.6 | 6.8 | 59.6 |
| Sham_M_65 | 93.6 | 0.3 | 1.8 | 66.8 |
| Sham_M_65* | 95.1 | 0.3 | 0.6 | 66.7 |
| Sham_P_23 | 88.6 | 3.1 | 3.2 | 48.6 |

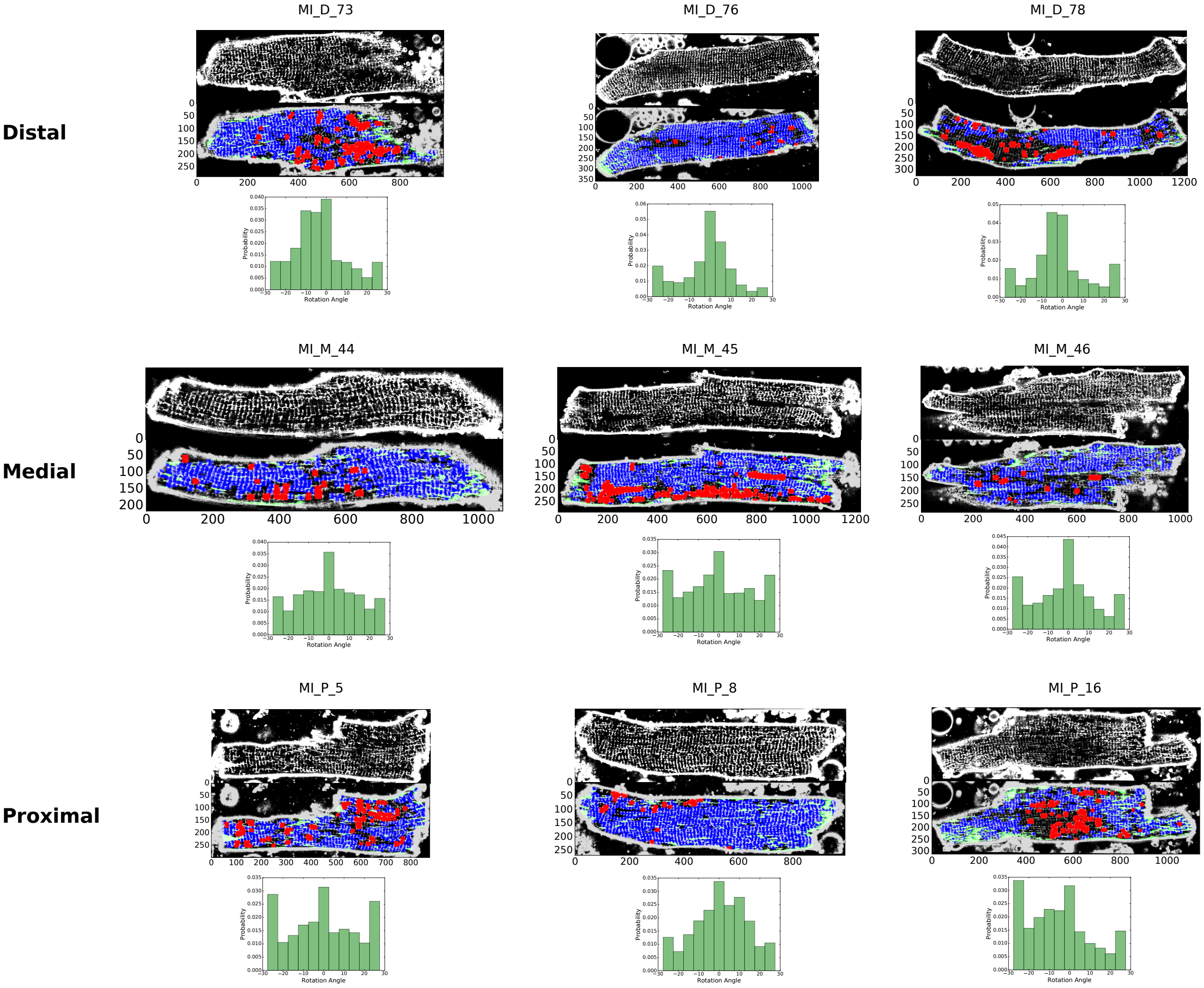


Figure S4: Results for all isolated infarct myocytes considered in this study. Isolated myocyte file name (top sub-row). Preprocessed image (second sub-row from top). Algorithm output marking transverse tubules, longitudinal tubules, and tubule absence in blue, green, and red, respectively (second sub-row from bottom). Histogram of transverse tubule detection angle (bottommost sub-row).

for each defined probability given in Table S8. This analysis was conducted using the matched filters provided herein extruded along the z-axis to form three dimensional filters. Due to increased signal integration, the SNR threshold for transverse tubule filtering had to be increased from 0.35 to 0.8 and longitudinal tubule filtering had to be increased from 0.6 to 0.9. Likewise, the standard deviation threshold for longitudinal filtering was removed and for tubule absence was increased from 0.1 to 0.5. Our analysis, shown in Table S8, indicates the MatchedMyo algorithm performs remarkably well with three dimensional classification with little optimization.

Table S6: Quantification of content present within myocytes from an infarcted rat model, * indicates the myocyte was analyzed with a mask for a region of interest excluding an assumed organelle. See S.3.1 for a description of tissue preparation. TT denotes a "hit" from the transverse tubule filter. Similarly, LT and TA denote "hits" from the longitudinal and tubule absence filters, respectively.

| Proximity | Myocyte | TT (%) | LT (%) | TA (%) | % WT Hits Within 5° of Minor Axis |
|---|---|---|---|---|---|
| Distal | MI_D_73 | 53.2 | 9.3 | 17.3 | 41.0 |
| | MI_D_76 | 75.6 | 6.4 | 2.6 | 54.5 |
| | MI_D_78 | 38.9 | 6.3 | 14.0 | 49.4 |
| | MI_D_78* | 47.9 | 2.6 | 8.7 | 46.5 |
| Intermediate | MI_M_44 | 62.5 | 9.7 | 064 | 36.1 |
| | MI_M_45 | 53.1 | 12.0 | 19.7 | 32.6 |
| | MI_M_45* | 59.5 | 7.2 | 23.4 | 31.8 |
| | MI_M_46 | 55.2 | 8.1 | 3.2 | 42.0 |
| Proximal | MI_P_5 | 50.0 | 9.1 | 17.7 | 31.2 |
| | MI_P_8 | 76.5 | 11.3 | 3.6 | 39.5 |
| | MI_P_16 | 38.7 | 15.8 | 13.4 | 32.4 |

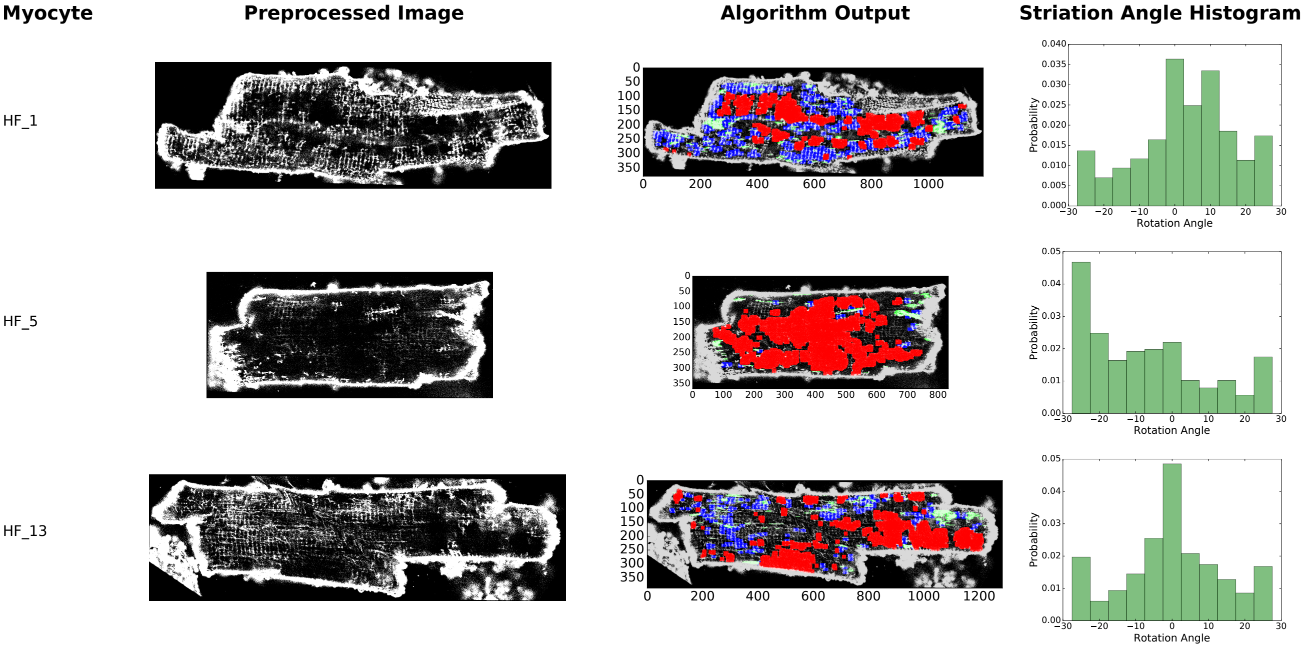


Figure S5: Results for all isolated myocytes from ascending aortic banded rat models considered in this study. Isolated myocyte file name (leftmost column). Preprocessed image (second column from left). Algorithm output marking transverse tubules, longitudinal tubules, and tubule absence in blue, green, and red, respectively (second column from right). Histogram of transverse tubule detection angle (rightmost column).

Table S7: Quantification of content present within myocytes from a ascending aortic banded rat model. TT denotes a "hit" from the transverse tubule filter. Similarly, LT and TA denote "hits" from the longitudinal and tubule absence filters, respectively.

| Myocyte | TT (%) | LT (%) | TA (%) | % TT Hits Within 5° of Minor Axis |
|---|---|---|---|---|
| HF_13 | 16.7 | 4.9 | 29.2 | 45.2 |
| HF_1 | 36.5 | 6.0 | 27.9 | 38.1 |
| HF_5 | 3.0 | 3.0 | 55.0 | 25.0 |

With further optimization, the minor overestimation of morphological content could be ameliorated.

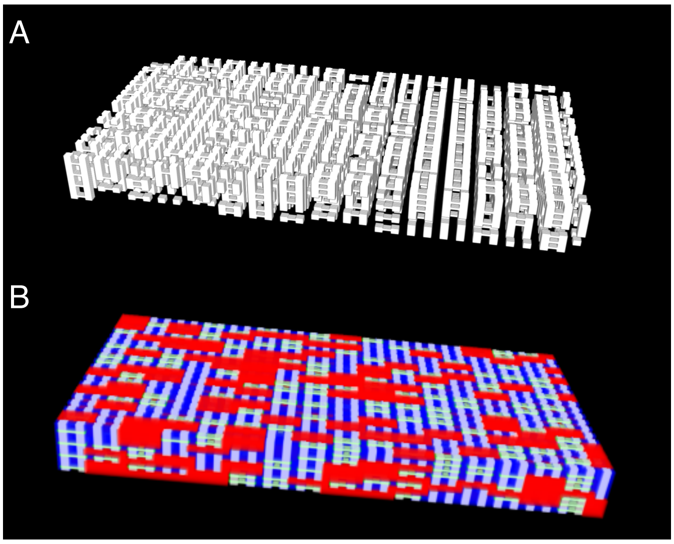


Figure S6: Proof of concept three dimensional classification on simulated data. (A) Sample simulated data and (B) resulting classification with TT (blue), LT (green), and TA (red) marked using the MatchedMyo algorithm.

| Unit Cell Probability (TT, LT, TA) | TT Content (Actual/Measured) | LT Content (Actual/Measured) | TA Content (Actual/Measured) |
|---|---|---|---|
| 0.55, 0.3, 0.15 | 0.255<br>0.267 ± 0.002 | 0.054<br>0.059 ± 0.001 | 0.15<br>0.232 ± 0.006 |
| 0.85, 0.0, 0.15 | 0.255<br>0.310 ± 0.003 | 0.0<br>0.0 | 0.15<br>0.233 ± 0.009 |
| 0.7, 0.15, 0.15 | 0.255<br>0.289 ± 0.002 | 0.027<br>0.030 ± 0.001 | 0.15<br>0.233 ± 0.006 |
| 0.7, 0.0, 0.3 | 0.21<br>0.241 ± 0.003 | 0.0<br>0.0 | 0.3<br>0.44 ± 0.01 |
| 1.0 , 0.0, 0.0 | 0.3<br>0.379 ± 0.0 | 0.0<br>0.0 | 0.0<br>0.0 |
| 0.55, 0.15, 0.3 | 0.21<br>0.219 ± 0.004 | 0.027<br>0.0284 ± 0.001 | 0.3<br>0.44 ± 0.01 |
| 0.85, 0.15, 0.0 | 0.3<br>0.359 ± 0.001 | 0.027<br>0.0299 ± 0.001 | 0.0<br>0.0 |
| 0.7, 0.3, 0.0 | 0.3<br>0.337 ± 0.001 | 0.054<br>0.059 ± 0.002 | 0.0<br>0.0 |
| 0.4, 0.3, 0.3 | 0.21<br>0.193 ± 0.003 | 0.054<br>0.057 ± 0.001 | 0.3<br>0.440 ± 0.008 |

Table S8: Results of extension of MatchedMyo algorithm to three dimensional classification of simulated cells. Simulated cell were generated based upon sequential addition of unit cells representing TT, LT, and TA morphology. Analysis consisted of classification of 20 replicates for each morphology probability combination.

#### S.4.3 Comparison of MatchedMyo performance to Fourier Transform analysis method

To ensure the accuracy of T-system classification by the MatchedMyo algorithm we compared our results with measures of TT density using a Fourier Transform method akin to the $TT_{Power}$ method developed by Wei et al [13].

**Method -** To measure T-system density using the Fourier Transform method, the following steps were taken:

1. Read in pre-oriented isolated cardiomyocyte confocal images with extracellular content and SL masked.
2. Smooth images with 3X3 kernel to attenuate noise.

3. Normalize image to sum of pixel intensities of image
4. Take Fourier Transform and take the log of the real component of the Fourier Transform to obtain the Power Spectral Density (PSD).
5. Pick out off-peak maximum of PSD as measure of TT density.

**Results -** The results of the analysis are shown in Table S9. Results of the analysis parallel results obtained from the MatchedMyo algorithm. Namely, sham models present the strongest measure of TT density, TT density increases as distance from infarct is increased, and TT density is the lowest in thoracic aortic banding (TAB) models.

| Case | Off-Peak PSD Value |
|---|---|
| Sham | -1.68 ± 0.44 |
| MI_D | -2.29 ± 0.10 |
| MI_M | -2.68 ± 0.09 |
| MI_P | -2.73 ± 0.21 |
| TAB | -2.99 ± 0.09 |

Table S9: Values obtained from analysis of TT density via Fourier Transform method. As with the MatchedMyo algorithm, TT density is strongest in sham models, TT density decreases as proximity to infarct increases, and TT density is the least in TAB models. Data presented as mean ± SD.

### S.5 SUPPLEMENTARY DISCUSSION

#### S.5.1 Limitations

While we demonstrate that matched filtering approach provides a step forward toward classification of microscopy data, several limitations must be considered. Brute force application of the approach to large images can quickly become cost-prohibitive, as the memory required to process large images scales rapidly with size. Hence, for characterization of large tissue sections, the image could be discretized into smaller sub-images, or downsampled via wavelet analysis [14] or eigen-analysis [15] to reduce the information density. Additionally, application of the algorithm to diverse multi-cellular microscopy images would require the user to tune the threshold parameters to optimize performance, given variations in immunofluorescence labeling, microscope performance, and cell structure. In this study, we determined optimal parameters through ROC analysis, yet this approach still necessitated hand annotation to produce training data. In contrast, detection algorithms such as AUTOTT [16] have relatively few parameters requiring optimization and thus may be less difficult to apply to new data sets. Lastly, for filters that have considerable spectral information in common, it can be challenging to unambiguously discriminate between similar data features. In this study, we considered filters with strongly different features, e.g. the longitudinal features were orthogonal to the transverse, which reduced overlap. Nevertheless, we found that introducing a penalty parameter was necessary to improve detection performance (see Sect. S.2.3 for more details).

### S.6 SUPPLEMENTARY FIGURES

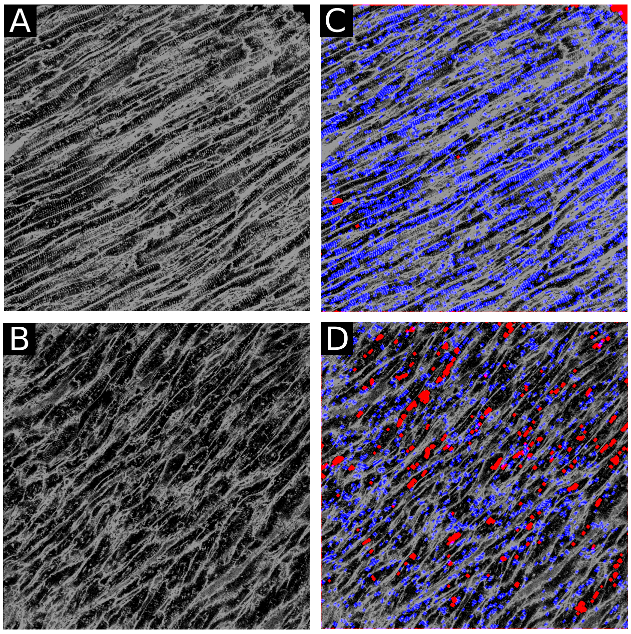


Figure S7: Application of TT filter (blue) and TA filter (red) to 3.9x4.1 $mm^2$ sections of infarcted rabbit tissue. (A) and (B) confocal images from regions distal and proximal to an infarct, respectively. (C) and (D) algorithm output using TT and TA filter. Striations arise from close apposition of punctate TT patterns typically observed in rabbit cardiomyocytes [2]. Relative to the distal region, the proximal region exhibited 45.8% as much TT content. Additionally, the proximal region exhibited 574% more T-system absence than the distal region.

[1] Z. Fegyver. *Calculate Standard Deviation: Case of Sliding Window.*

[8] T. Seidel, A. Sankarankutty, and F. Sachse. "Remodeling of the transverse tubular system after myocardial infarction in rabbit correlates with local fibrosis: A potential role of biomechanics". In: *Progress in Biophysics and Molecular Biology* (July 2017). ISSN: 00796107.

[14] G. Strang and T. Nguyen. *Wavelets and Filter Banks.* 1st ed. Wellesley-Cambridge Press, 1996. ISBN: 9780961408879.

[15] M. Turk and A. Pentland. "Eigenfaces for Recognition". In: *Journal of Cognitive Neuroscience* 3.1 (Jan. 1991), pp. 71–86. ISSN: 0898-929X.

[16] A. Guo and L.-S. Song. "AutoTT: automated detection and analysis of T-tubule architecture in cardiomyocytes." In: *Biophysical journal* 106.12 (June 2014), pp. 2729–36. ISSN: 1542-0086.
